# Data-driven brain-types and their cognitive consequences

**DOI:** 10.1101/237859

**Authors:** Joe Bathelt, Amy Johnson, Mengya Zhang, the CALM team, Duncan E. Astle

## Abstract

The canonical approach to exploring brain-behaviour relationships is to group individuals according to a phenotype of interest, and then explore the neural correlates of this grouping. A limitation of this approach is that multiple aetiological pathways could result in a similar phenotype, so the role of any one brain mechanism may be substantially underestimated. Building on advances in network analysis, we used a data-driven community-clustering algorithm to identify robust subgroups based on white-matter microstructure in childhood and adolescence (total N=313, mean age: 11.24 years). The algorithm indicated the presence of two equal-size groups that show a critical difference in FA of the left and right cingulum. These different ‘brain types’ had profoundly different cognitive abilities with higher performance in the higher FA group. Further, a connectomics analysis indicated reduced structural connectivity in the low FA subgroup that was strongly related to reduced functional activation of the default mode network.

**Graphical abstract:** 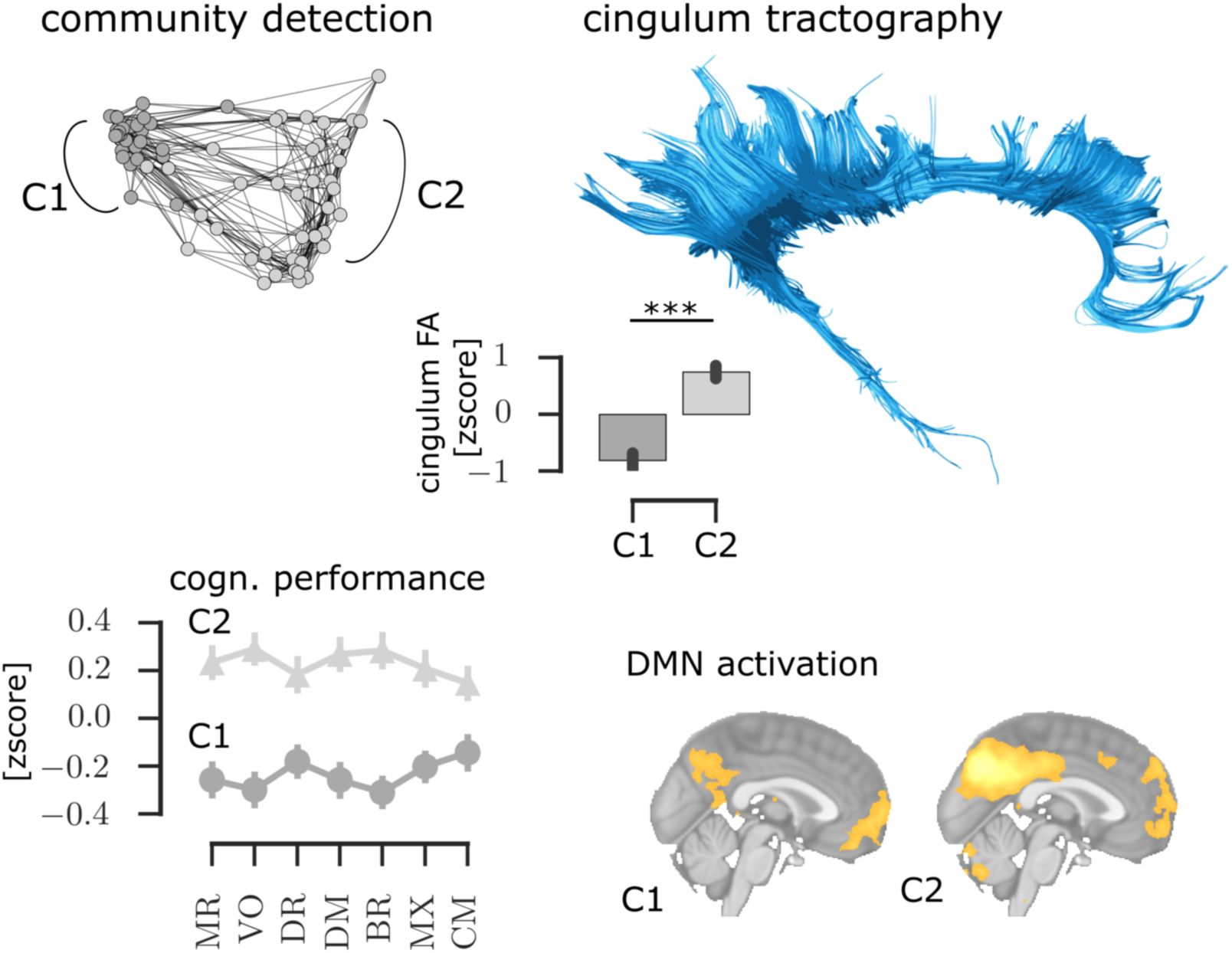

## Introduction

Differential psychology is an influential strand of modern psychology, concerned with identifying dimensions upon which individuals differ. This approach has been applied at different points across development, from childhood to old adulthood, and has been central to our understanding of typical and atypical behaviour and psychopathology (Lubinski 2000; Cronbach 1957). Our understanding of the brain basis mechanisms associated with these differences is based almost entirely on mapping them via correlations with brain differences. This has established many consistent and significant brain-behaviour relationships in health and disease. But understanding neurobiology only through our prior understanding of cognition or behaviour has drawbacks. Firstly, our understanding of the neurobiology is entirely constrained by the choice of cognitive measures. Secondly, brain-behaviour relationships established using this logic are difficult to replicate (Poldrack and Yarkoni 2016; Uttal 2001). This partly reflects this dependency on task selection, which differs across research groups and may only partially tap the dimension of interest. But more critically, multiple aetiological pathways could lead to disorders with superficially similar phenotypes (Stevens et al. 2017; Jones et al. 2014; Fried and Kievit 2015). In short, grouping purely by cognitive or behavioural phenotype is no guarantee of common underlying neurobiology.

At present, there are few alternatives to complement this standard approach. The current study shows that it is possible to group individuals by brain organization itself, rather than by cognitive or behavioural phenotype. That is, we identified stable subgroups of individuals with similarly organized brains, which are distinct from members of other subgroups. In doing so we hoped to identify the elements of neurobiology most critical for dividing individuals into their respective subgroups and their wider functional consequences. Identifying mechanisms of brain difference a priori has the potential to highlight key organizational principles that we may never capture by only looking for neural correlates of messy pre-defined phenotypes. This was made possible by combining recent advances in network analysis and community-clustering algorithms, with measures of microstructural white matter organization.

## The importance of white matter in cognitive development

White matter makes up around half of the human brain and plays a critical role as the main conductor of neural signalling. White matter also shows prolonged postnatal changes that extend into the third decade of life (Lebel, Treit, and Beaulieu 2017) with pronounced development during mid-childhood and adolescence (Westlye et al. 2009). In humans, the method of choice for measuring differences in white matter *in vivo* is diffusion-weighted imaging (DWI). It quantifies the degree to which diffusion of water molecules is restricted by the tightly-packed parallel axons that make up white matter. Differences in DWI-derived measures have been found to relate to individual differences across a range of cognitive domains. For instance, language processing is related to the maturation of the arcuate fasciculus, a white matter tract that connects the frontal and superior temporal lobe (Skeide, Brauer, and Friederici 2015); working memory performance is associated with the integrity of the superior longitudinal fasciculus, a tract connecting frontal and parietal regions (Burzynska et al. 2011); executive control is linked to the integrity of frontal and parietal connections alongside additional connections to motor control regions (Chaddock-Heyman et al. 2013). Differences in white matter organisation have also been identified in various neurodevelopmental disorders. For instance, dyslexia is associated with a reduced organisation in white matter pathways along the left dorsal and ventral language pathways (Zhao et al. 2016); children with Attention Deficit Hyperactivity Disorder (ADHD) show reduced white matter organisation of the corpus callosum and major tracts of the right hemisphere (Wu et al. 2016); and reduced integrity of connections between the limbic system and frontal and temporal cortex has been reported for autism spectrum disorder (ASD) (Ameis and Catani 2015). In summary, DWI measures of white matter organisation are sensitive to typical variation in cognitive abilities and show differences in common neurodevelopmental disorders.

## Using community detection to identify subgroups with similar white matter organisation

Network science is the study of complex networks, which represent relationships between data as a network of nodes connected by edges. This methodological approach provides a mathematical tool for quantifying the organisation of networks and the relationships between the nodes within them (Bullmore and Sporns 2009). Defining subdivisions of highly-connected nodes within a network, so-called communities, is an area of network science that has received considerable attention as it applies to many real-world problems (Barabasi 2016). In our case, the network represents the similarity of white matter organization in 20 major tracts cross individuals. Community detection makes it possible to define subgroups of participants that are most similar while being as distinct as possible from other subgroups. This approach has been successfully applied to distinguish differences in behaviour within heterogeneous groups, such as subgroups of neuropsychological function in typically-developing children (Fair et al. 2012) and subgroups of executive function-related behavioural problems in children who struggle in school (Bathelt, Holmes, and Astle 2017). In summary, community-clustering provides an ideal method to identify subgroups of individuals with similar characteristics.

## Participants & Methods Participants

### Participants

The participants were drawn from three separate studies of development. The participants included in each study are characterised in the following sections

#### The Nathan-Kline Institute Rockland Sample (NKI-RS)

The enhanced Nathan Kline Institute-Rockland Sample (NKI-RS) is an ongoing, institutionally-centred endeavour aimed at creating a large-scale community sample of participants across the lifespan. Details about the sample are described on the NKI website (http://fcon_1000.projects.nitrc.org/indi/enhanced/). NKI sample data is available to researchers upon request. For the current study, data from all participants who had structural imaging (T1, diffusion-weighted) and age information available was requested (date of data access: 18 July 2017). The study was approved by the NKI institutional review board and all adult and child subjects provided informed consent (Nooner et al. 2012)

#### Centre for Attention, Learning, and Memory (CALM)

For this sample, children aged between 5 and 18 years were recruited on the basis of ongoing problems in attention, learning, language and memory, as identified by professionals working in schools or specialist children’s services in the community. Following an initial referral, the CALM staff then contacted referrers to discuss the nature of the children’s problems. If difficulties in one or more areas of attention, learning, language or memory were indicated by the referrer, the family were invited to the CALM clinic at the MRC Cognition and Brain Sciences Unit in Cambridge for a 3-hour assessment. This assessment included the cognitive assessments reported here. Exclusion criteria for referrals were significant or severe known problems in vision or hearing that were not corrected or having a native language other than English. Written parental consent was obtained and children provided verbal assent. Families were also invited to participate in MRI scanning on a separate visit. Participation in the MRI part of the study was optional and required separate parental consent and child assent. Contra-indications for MRI were metal implants, claustrophobia, or distress during a practice session with a realistic mock MRI scanner. This study was approved by the local NHS research ethics committee (Reference: 13/EE/0157).

#### Attention and Cognition in Education (ACE)

This sample was collected for a study investigating the neural, cognitive, and environmental markers of risk and resilience in children. Children between 7 and 12 years attending mainstream school in the UK, with normal or corrected-to-normal vision or hearing, and no history of brain injury were recruited via local schools and through advertisement in public places (childcare and community centres, libraries). Participating families were invited to the MRC Cognition and Brain Sciences Unit for a 2-hour assessment, which included the cognitive assessments reported here, and structural MRI scanning. Participants received monetary compensation for taking part in the study. This study was approved by the Psychology Research Ethics Committee at the University of Cambridge (Reference: Pre.2015.11). Parents provided written informed consent and children verbal assent.

### Cognitive assessments

#### Procedure

All children for whom we have cognitive data were tested on a one-to-one basis with a researcher in a dedicated child-friendly testing room at the MRC CBU. The battery included a wide range of standardized assessments of learning and cognition. Regular breaks were included throughout the session. Testing was split into two sessions for children who struggled to complete the assessments in one sitting. Measures relating a cognitive performance across different domains are included in this analysis. Tasks that were based on reaction times were not included in this analysis due to their different psychometric properties compared to the included tasks that were based on performance measures.

#### Fluid intelligence

Fluid intelligence was assessed on the Reasoning task of the Wechsler Abbreviated Scale of Intelligence, 2^nd^ edition (Wechsler 2011). Both children in the CALM and ACE sample completed this assessment.

#### Working Memory

The Digit Recall, Backward Digit Recall, Dot Matrix, and Mr X task of the Automatic Working Memory Assessment (AWMA) (Alloway et al. 2008) were administered individually. In Digit Recall, children repeat sequences of single-digit numbers presented in an audio format. In Backward Digit Recall, children repeat the sequence in backwards order. These tasks were selected to engage verbal short-term and working memory, respectively. For the Dot Matrix task, the child was shown the position of a red dot for 2 seconds in a series of four by four matrices and had to recall this position by tapping the squares on the computer screen. In the Mr X task, the child retains spatial locations whilst performing interleaved mental rotation decisions. These tasks were selected to engage visual short-term and working memory, respectively. These assessments were the same in the CALM and ACE sample.

#### Vocabulary

For the CALM sample, the Peabody Picture Vocabulary Test, Fourth Edition (PPVT-4) (Dunn and Dunn 2007) was used to assess receptive vocabulary knowledge. Children were required to select one of four pictures showing the meaning of a spoken word. In the ACE sample, the Vocabulary subtest of the Wechsler Abbreviated Scale of Intelligence, 2nd edition (Wechsler 2011), was used. For this task, children had to define words that were presented verbally and visually, and correct definitions were scored.

#### Long-term memory

For the CALM sample, the Stories task of the Children’s Memory Scale (Cohen 1997) was used to assess long-term memory. Children were read two short stories and were asked to recall these stories after a delay of 10 min. No long-term memory task was used in the ACE sample.

### MRI data acquisition

#### NKI-RS

Subjects in the NKI sample underwent a scan session using a Siemens TrioTM 3.0 T MRI scanner. T1-weighted images were acquired a magnetization-prepared rapid gradient echo (MPRAGE) sequence with 1mm isotropic resolution. Diffusion scans were acquired with an isotopic set of gradients with 64 directions using a weighting factor of b=1000s*mm^−2^ and an isotropic resolution of 2mm. Details about the scan sequences are described elsewhere (http://fcon_1000.projects.nitrc.org/indi/enhanced/mri_protocol.html).

#### CALM & ACE

Magnetic resonance imaging data were acquired at the MRC Cognition and Brain Sciences Unit, Cambridge U.K. All scans were obtained on the Siemens 3 T Tim Trio system (Siemens Healthcare, Erlangen, Germany), using a 32-channel quadrature head coil. For ACE, the imaging protocol consisted of two sequences: T1-weighted MRI and a diffusion-weighted sequence. For CALM, the imaging protocol included an additional resting-state functional MRI (rs-fMRI) sequence. T1-weighted volume scans were acquired using a whole brain coverage 3D Magnetisation Prepared Rapid Acquisition Gradient Echo (MP-RAGE) sequence acquired using 1mm isometric image resolution. Echo time was 2.98 ms, and repetition time was 2250 ms. Diffusion scans were acquired using echo-planar diffusion-weighted images with a set of 60 non-collinear directions, using a weighting factor of b=1000s*mm-2, interleaved with a T2-weighted (b=0) volume. Whole brain coverage was obtained with 60 contiguous axial slices and isometric image resolution of 2mm. Echo time was 90 ms and repetition time was 8400 ms. For functional scans, a total of 270 T2*-weighted whole-brain echo planar images (EPIs) were acquired (time repetition[TR] = 2 s; time echo [TE] = 30 ms; flip angle = 78 degrees, 3×3×3mm). During this scan, participants were instructed to lie still with their eyes closed and not fall asleep.

#### MRI quality control

Participant movement may significantly affect the quality of MRI data and may bias statistical comparisons. Several steps were taken to assure good MRI data quality and minimize potential biases of participant movement. First, children were instructed to lie still and were trained to do so in a realistic mock scanner prior to the actual scan. Second, all T1-weighted images and FA maps were visually inspected by a trained researcher (J.B.) to remove low quality scans. Further, the quality of the diffusion-weighted and resting-state data were assessed by calculating the displacement between subsequent volumes in the sequence. Only dwi data with between-volume displacement below 3mm and resting-state data with displacement below 0.5mm were included in the analysis (see Table 1 for attainment).

**Table 1:** Overview of sample characteristics in the NKI, ACE, and CALM sample

|  | n: useable MRI<br>(total) | Gender:<br>male-female | Age: mean (SD) | Age range |
| --- | --- | --- | --- | --- |
| NKI | 74 (117) | 43/31 | 13.93 (3.164) | 6.97-21.58 |
| ACE | 74 (86) | 35/39 | 9.99 (1.525) | 6.92-12.65 |
| CALM | 165 (206) | 109/56 | 9.81 (1.191) | 5.92-17.92 |

## Microstructural integrity of major white matter tracts

This analysis aimed to identify white matter tracts that show robust inter-individual differences during development (see Figure 1A). Diffusion-weighted images were pre-processed to create a brain mask based on the b0-weighted image (FSL BET) (Smith 2002) and to correct for movement and eddy current-induced distortions (*eddy*) (Graham, Drobnjak, and Zhang 2016). Subsequently, the diffusion tensor model was fitted and fractional anisotropy (FA) maps were calculated (*dtifit*). Images with a between-image displacement great than 3mm as indicated by FSL eddy were excluded from further analysis. All steps were carried out with FSL v5.0.9 and were implemented in a pipeline using NiPyPe v0.13.0 (Gorgolewski et al. 2011). To extract FA values for major white matter tracts, FA images were registered to the FMRIB58 FA template in MNI space using symmetric diffeomorphic image registration (*SyN*) as implemented in ANTS v1.9 (Avants et al. 2008). Visual inspection indicated good image registration for all participants. Subsequently, binary masks from a white matter atlas in MNI space (Mori, Oishi, and Faria 2009) were applied to extract FA values for 20 major white matter tracts. An additional analysis was carried out with Generalized FA (GFA, Tuch 2004) based on a constant solid angle model (details see connectome processing) to account for crossing fibres.

**Figure 1A.**
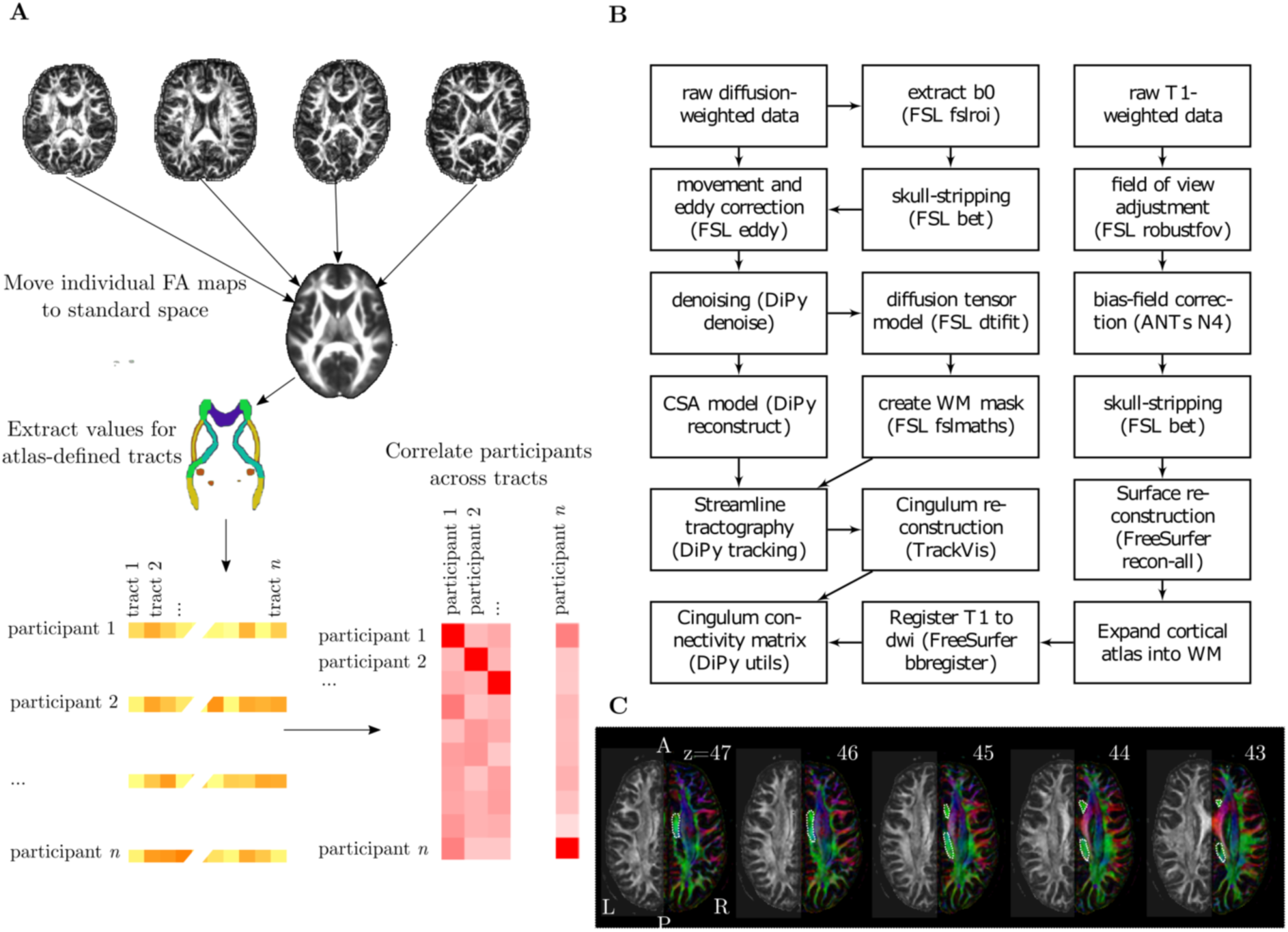
Illustration of processing steps to obtain estimates of similarity between individuals across major white matter tracts **B** Overview of processing steps to derive structural connectivity of the cingulum **C** Illustration of the ROI used for tractography of the cingulum. The ROI is marked with a dotted line on consecutive FA maps coloured according to the direction of diffusion. The left side of each figure shows the FA values.

## Community detection

Community detection is an optimisation clustering method. Networks in the current analysis represented the child-by-child correlations across the FA values in 20 white matter tracts. The community algorithm starts with each network node, i.e. child, in a separate community and then iteratively parcellates the network into communities to increase the quality index (Q), which represents the segration between communities, until a maximum is reached. The current study used the algorithm described by Rubinov and Sporns (Rubinov and Sporns 2010) as implemented in the Brain Connectivity Toolbox (https://sites.google.com/site/bctnet/) version of August 2017, which is an extension of the method described by Blondel et al. (Blondel et al. 2008) to networks with positive and negative edges. This algorithm is not deterministic and may yield different solutions at each run. In order to reach a stable community assignment, we applied the consensus clustering method described by Lancichinetti and Fortunato (Lancichinetti and Fortunato 2012). In short, an average community assignment over 100 iterations was generated. The community assignment was then repeated further until the community assignment did not change between successive iterations.

In order to test the reliability of the community detection algorithm under varying conditions, random networks with known community structure were created. The networks consisted of 100 nodes with 4 modules. The connection likelihood within and between clusters was systematically varied between 0.1 and 0.9. The quality index of the community structure was calculated at each combination of between- and within-cluster connection likelihood. The results indicated a high-quality index for networks with higher within-cluster than outside-cluster connection likelihood (see Figure 2A). High connection density outside of clusters had a large influence, even when the connection likelihood within modules was very high. We further tested the robustness of the community assignment by adding increasing percentages of random Gaussian noise (mu=0, sigma=1) to the network matrix and repeated the consensus clustering procedure (see Figure 2B). The quality index indicated a good separation of the clusters between 5 and 30% noise (Q between 0.62 and 0.65). No stable assignment could be reached at 35% of noise and above. In short, these results indicate that the community assignment is robust to a considerable amount of noise. Further, the Keringhan-Lin algorithm was used to test the grouping with an alternative method that can integrate negative edge weights (Kernighan and Lin 1970). The results indicated nearly identical results with this alternative method (see Results section).

**Figure 2.**
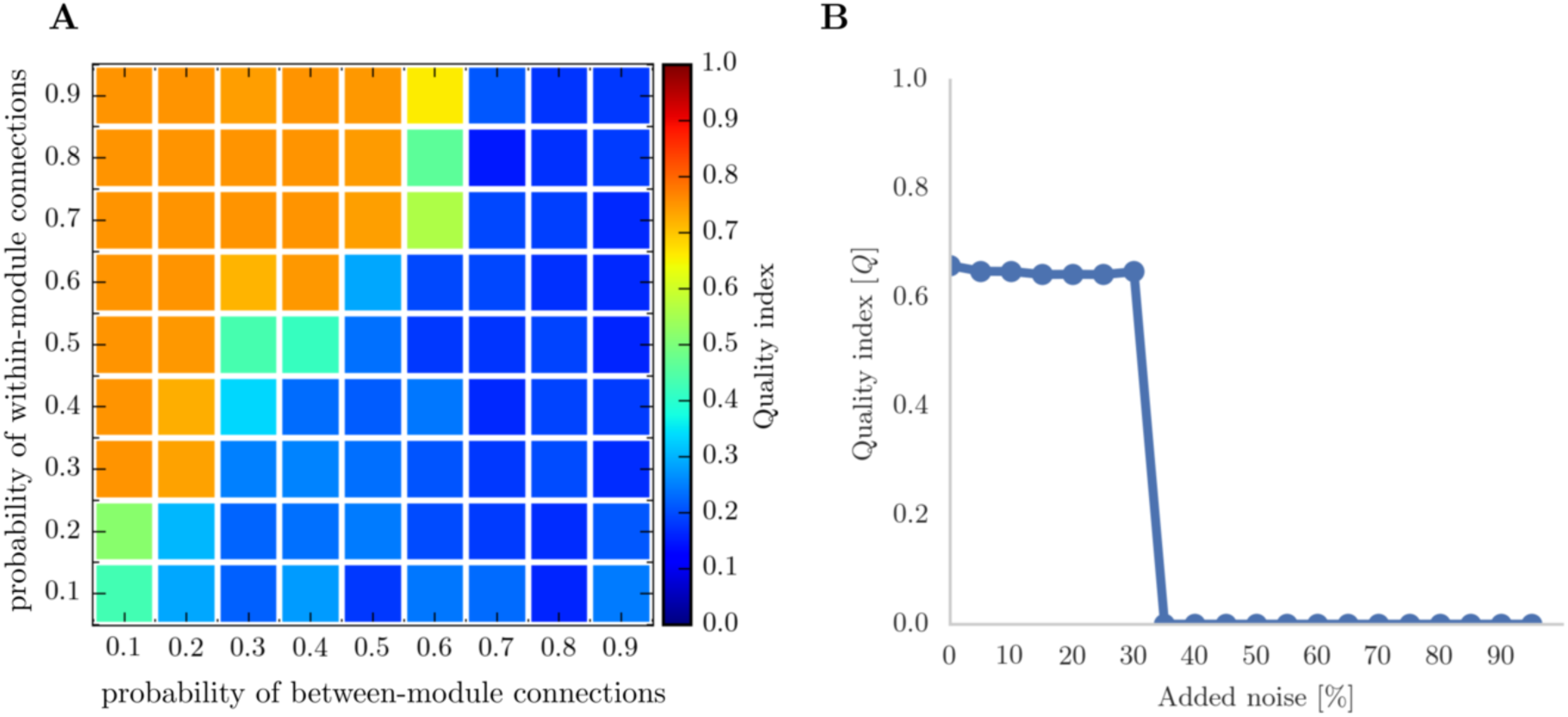
Results of robustness testing using networks with a known community structure. A Quality index in networks with different probabilities of within- and between-community connections **B** Quality index in networks with increasing ratios of Gaussian noise.

In order to test if the clustering identified in one sample generalized to a sample that was not used to inform the algorithm, we compared the within versus-between-cluster summed correlation to a permutation sample with random cluster assignment over 1000 repetitions. If the clustering provides a good account of the data, then the summed within-cluster correlation should be considerably higher than the between-sample correlation and both should be significantly different from random cluster assignments. This provides a good way of validating the initial community clustering solution with the subsequent independent datasets.

We further tested the generalizability of the prediction for behavioural performance based on brain type assignment. For this purpose, we randomly split the data into a training (ACE: 60%, CALM: 80%) and a test set (ACE: 40%, CALM: 20%). In the training set, we determined the optimal percentile cut-off of behavioural scores to maximize the prediction accuracy of the brain type grouping. Then, we tested the accuracy of the behavioral prediction in the hold-out test set using the percentile threshold identified in the training set.

## Tractography of the cingulum

To foreshadow our results, the clustering algorithm identified the cingulum as critical for determining subgroup membership. We subsequently followed this up with tractography. For the tractography, a virtual in-vivo dissection approach was applied (Catani and de Schotten 2008) (see Figure 1B for an overview of the processing steps). For this purpose, MRI scans were converted from the native DICOM to compressed NIfTI-1 format using the dcm2nii tool developed at the McCauseland Centre for Neuroimaging ([http://www.mccauslandcenter.sc.edu/mricro/mricron/dcm2nii.html]). Subsequently, the images were submitted to the DiPy v0.8.0 implementation (Garyfallidis et al. 2014) of a non-local means de-noising algorithm (Coupe et al. 2008) to boost signal-to-noise ratio. Next, a brain mask of the b0 image was created using the brain extraction tool (BET) of the FMRIB Software Library (FSL) v5.0.8. Motion and eddy current correction was applied to the masked images using FSL routines. Finally, fractional anisotropy maps were created based on a diffusion tensor model fitted through the FSL *dtifit* algorithm (Behrens et al. 2003). A constant solid angle (CSA) was fitted to the 60-gradient-direction diffusion-weighted images using a maximum harmonic order of 8 using DiPy. Next, probabilistic whole-brain tractography was performed based on the CSA model with 8 seeds in any voxel with a General FA value higher than 0.1. The step size was set to 0.5 and the maximum number of crossing fibres per voxel to 2. The cingulum in the left and right hemisphere was reconstructed in the native space of each participant by drawing a single ROI on 10-15 consecutive axial slices that followed the anatomy of the cingulum as described in (Catani and de Schotten 2008) (see Figure1C). To check the reliability of the cingulum tractography, two researchers performed tractography independently on 40 datasets. The spatial correlation between the density maps indicated very good inter-rater agreement (left: mean=0.95, SE=0.005, range=0.86-0.990; right: mean=0.95, SE=0.005, range=0.87-0.990).

## Structural connectivity of the cingulum

To estimate the structural connectivity of the cingulum, streamlines included in the cingulum tractography that intersected with cortical and subcortical ROIs were counted. For ROI definition, T1-weighted images were preprocessed by adjusting the field of view using FSL’s *robustfov*, non-local means denoising in DiPy, deriving a robust brain mask using the brain extraction algorithm of the Advanced Normalization Tools (ANTs) v1.9 (Avants et al. 2011), and submitting the images to recon-all pipeline in FreeSurfer v5.3 (http://surfer.nmr.mgh.harvard.edu). Regions of interests (ROIs) were based on the Desikan-Killiany parcellation of the MNI template (Desikan et al. 2006) with 34 cortical ROIs per hemisphere and 17 subcortical ROIs (brain stem, and bilateral cerebellum, thalamus, caudate, putamen, pallidum, hippocampus, amygdala, nucleus accumbens). The surface parcellation of the cortex was transformed to a volume using the aparc2aseg tool in FreeSurfer. Further, the cortical parcellation was expanded by 2mm into the subcortical white matter using in-house software. In order to move the parcellation into diffusion space, a transformation based on the T1-weighted volume and the b0-weighted image of the diffusion sequence was calculated using FreeSurfer’s *bbregister* and applied to volume parcellation. For each pairwise combination of ROIs, the number of streamlines intersecting both ROIs was estimated and transformed into a density map. The intersection was symmetrically defined, i.e. streamlines starting and ending in each ROI were averaged.

In order to minimize false positive connections, the streamline density matrices were thresholded so that only connections with more than 5 streamlines were retained. Further, the matrices were consensus-thresholded to only contain connections that were found in at least 60% of the sample (de Reus et al., 2009). Subsequently, the streamline density matrices were log_10_-scaled.

Individual connections that consistently differed between the groups were identified through hold-out cross-validation in a dataset combining tractography results from the ACE and CALM sample (n=239). The dataset was randomly split into a training (80%, n=188) and a test set (20%, n=47). Within the training set, connections that showed a significant difference between cingulum-defined groups on a Mann-Whitney-U test that were consistently observed across 4 random split of the training data (n=47) were selected. Subsequently, differences between groups in the strength of the selected connections were checked in the hold-out test set. Only connections that showed consistent differences in the training and test set are presented in the Results section.

## Default mode network activation

Resting-state functional MRI data were processed to obtain functional activation within the default mode network. Only participants who completed the full resting-state sequence had full coverage of the brain in the resting-state sequence, and also had useable T1-weighted data were included in the analysis. For motion and eddy current correction, volumes were co-registered to the middle volume using *mcflirt* (Jenkinson et al. 2002). For quality control, the frame-wise displacement was calculated as the weighted average of the rotation and translation parameters (Power et al. 2012) using *fsl_motion_outliers*. Participants with a frame-wise displacement above 0.5 on this measure were excluded from the analysis. Complete data was available for 124 participants (78 male, Age: mean=10.00, SE=0.196, group1: n=59, group2: n=58, no group: n=6). There was no significant difference in age between the sample included in the rsfMRI analysis and the sample included in the main analysis (t(284)=-0.74, p=0.458). The samples did not differ on any of the cognitive measures (MR: t=-0.85, *p*=0.396, Vocab: t=-0.80, *p*=0.423; DR: t=-0.73, *p*=0.465; DM: t=-0.62, *p*=0.539; BR: t=-0.80, *p*=0.427; MX: t=-0.23, *p*=0.820; CM: t=-0.74, *p*=0.461). Regarding data quality, there was no difference in frame-wise displacement between the groups defined through data-driven clustering (group 1: mean=0.01, SE=0.001; group2: mean=0.01, SE=0.004, t(122)=-0.71, p=0.480). The groups also did not differ in the number of motion outliers (group 1: mean=18.05, SE=1.007; group 2: mean=17.08, SE=1.154, t(122)=0.63, p=0.529). There was no significant correlation between the framewise displacement and any of the cognitive measures (Pearson correlation, MR: r=-0.12, *p*=0.188; Vocab: r=0.09, *p*=0.349; DR: r=-0.00, *p*=0.986; DM: r=-0.11, *p*=0.226; BR: r=0.07, *p*=0.456; MX: r=0.05, *p*=0.570; CM: r=0.11, *p*=0.207).

Functional connectivity was assessed using independent component analysis by means of the multivariate exploratory linear decomposition into a fixed set of 25 independent components (MELODIC; (Beckmann et al. 2005; Beckmann and Smith 2004)). The correspondence between the independent components derived in the data and the canonical networks described by Yeo et al. 2011 (Yeo et al. 2011) was established by calculating the spatial correlation between the maps. Two maps showed a high spatial correlation with the canonical default mode network (IC3: r=0.303, IC9: r=0.493). Two additional component maps also had a high spatial correlation with the default mode network map (r=0.286, r=0.276), but showed activation within the white matter and CSF and were therefore dismissed as artefacts. The default mode network in individual participants was calculated as described by Supekar and colleagues (Supekar et al. 2010). In brief, ICA with automatic dimensionality detection was performed for each participant. The groups did not differ in the number of ICA components generated (C1: mean=100.70, SE=1.821; C2: mean=102.19, SE=1.862; U=1549, *p*=0.254). Subsequently, a goodness-of-fit measure was calculated based on the average z-scored activation within the DMN-mask minus the activation outside of the DMN-mask and the component with the largest score was selected. To compare the spatial extent of activation within the individual DMN components between the groups, a permutation procedure with 5,000 repetitions and cluster-free threshold enhancement for correction of multiple comparisons as implemented in FSL randomise was used. To compare functional connectivity between groups, the average bandpass-filtered (0.01-0.1Hz) signal for a 8mm-sphere placed on the posterior cingulate cortex (PCC, MNI: 0,-52,18), medial prefrontal cortex (mPFC, MNI: 1, 50, −5), and the left and right temporoparietal junction (TPJ, MNI: −46, −68, 32; 46, −68, 32) and the partial correlation between mPFC and PCC was calculated controlling for the signals in the left and right TPJ (Heuvel et al. 2008).

## Statistical analysis

Shapiro-Wilk tests were used to test if data were normally distributed. If a significant deviation from normality was indicated (*p*>0.05), non-parametric tests were used. Specifically, Mann-Whitney U tests were used instead of independent t-tests. The Bonferroni method was used to correct for multiple comparisons unless otherwise stated.

## Results

The analysis followed a three-step sequential logic:

- The first aim was to determine subgroups of maximally similar white matter organisation using community detection in a population-representative sample of typical development (NKI sample).
- The second aim was to investigate potential cognitive correlates of white matter tracts that distinguished groups identified in the first part of the analysis. This was carried in large developmental samples that had detailed cognitive assessments and showed large variation in cognitive abilities (ACE & CALM sample).
- The third aim was to investigate the relationship between white matter variation in more detail by mapping the particular connections related to different aspects of cognitive ability and relating variation in white matter microstructure to functional connectivity in a sample with detailed cognitive assessments and multimodal neuroimaging data (ACE & CALM sample).

## Community clustering indicates the presence of two groups that differ in FA of the cingulum

Grouping children in the NKI sample according to their similarity of FA values within major white matter tracts using consensus clustering indicated the presence of two groups. The quality index indicated a good separation between groups (Q=0.47, Figure 3 A and B). The groups defined through community clustering did not differ in age (C1: n=34, mean=13.568, SE=0.59; C2: n=40, mean=14.23, SE=0.470; t-test: t(72)=-0.89, p=0.375; Equivalence test: t_lower_(65.94)=1.355, *p*=0.09, t_upper_(65.94), *p*=0.001 with δ=1.684) and did also not differ from the whole sample in proportion of males and females (C1: 24/10 male/female, χ^2^=0.06, *p*=0.800; C2: 21/19, χ^2^=0.05, *p*=0.825). Comparison of group assignment with an alternative clustering method (Kernighan-Lin algorithm) indicated very high agreement with an equal number of clusters using both methods and only 6 out 74 children being assigned to a different cluster depending on the algorithm.

**Figure 3.**
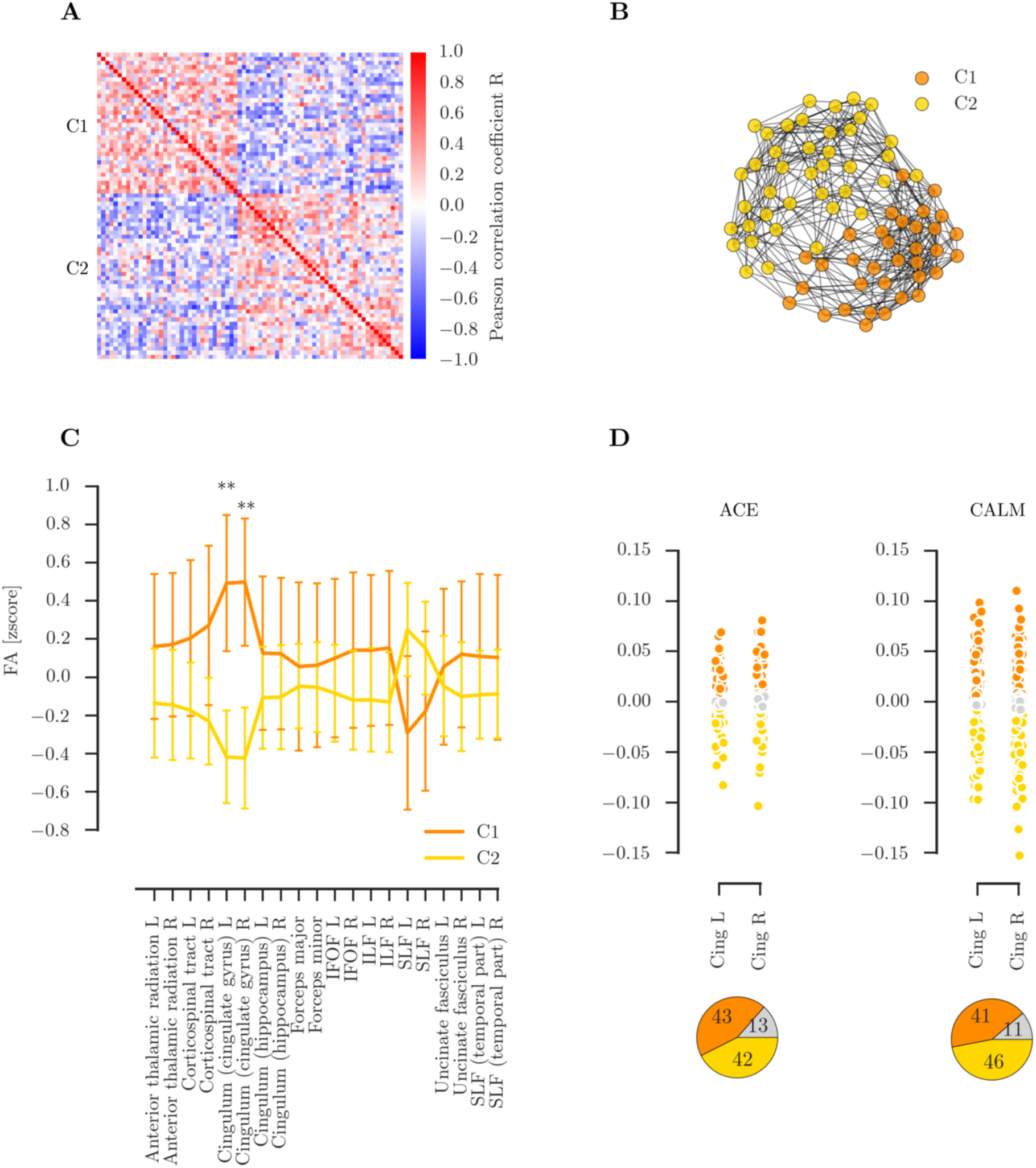
Overview of clustering based on fractional anisotropy (FA) in major white matter tracts in the NKI sample. **A**: Correlation within groups defined through community-based clustering of FA values in the NKI sample showing two distinct groups with high correlation within the cluster. **B**: Spring-layout depiction of the clusters identified through community clustering show good separation between the clusters (thresholded at R0.3 visualisation purposes). **C**: Differences in FA for each major white matter tract between the clusters identified through community clustering. The error bars show one standard error, the middle of the bar indicates the median. Significant differences between clusters were found for the left and right anterior Cingulum. **D**: FA values in the left and right anterior cingulum in the ACE and CALM with grouping according to the ranges identified in the NKI sample clusters. The bottom figure show the percentage of participants in each cluster range with orange for C1, yellow for C2, and grey for participants that fall outside of both ranges. Abbreviations: IFOF: inferior fronto-occipital fasciculus, ILF: inferior longitudinal fasciculus, SLF: superior longitudinal fasciculus

Next, FA values for all tracts were compared between the groups defined through community clustering. The groups differed on the FA of the left and right anterior cingulum (cingulate gyrus region) (left anterior cingulum: C1: mean=0.49, SE=0.179, C2: mean=-0.42, SE=0.121, Man-Whitney: U=304, *p*<0.001, *p*_corrected_<0.001; right anterior cingulum: mean=0.50, SE=0.167, C2: mean=-0.42, SE=0.132 Man-Whitney: U=300, *p*<0.001, *p*_corrected_<0.001). There were no significant differences between the groups for any other tract (all other pcorrected>0.1, see Figure 3C). To establish the potential influence of crossing fibres, we repeated the analysis with GFA values. This comparison indicated a significantly lower GFA in C2 for the left and right anterior cingulum (left: C1: mean=0.20, SE=0.004, C2: mean=0.19, SE=0.003, U=426, *p*=0.003, *p*_corrected_=0.03, right: C1: mean=0.22, SE=0.004, C2: mean=0.21, SE=0.003, U=430, *p*=0.003, *p*_corrected_=0.034). There were no significant differences in GFA for any other tract (all other *p*_corrected_>0.1).

The range of age-regressed, *z*-transformed values for the left and right cingulum was used to group children in the CALM and ACE sample (see Figure 3D). A majority of the samples fell within the range for C1 or C2 (ACE: C1: n=36 (42%), C2: n=43 (51%), not categorized: n=5 (6%); CALM: C1: n=95 (49%), C2: n=87 (45%), not categorized: n=11 (6%)). The proportion of participants being assigned to C1 or C2 was equal (ACE: *χ* ^2^=0.64, *p*=0.423; CALM: *χ*^2^=0.01, *p*=0.943). Comparison of the within-group versus between-group connection strength indicated that the grouping based on the clusters identified in the NKI sample provided a good account of the data for both the ACE and the CALM sample that significantly differed from random group assignments (ACE: intra-cluster= 64.85, *p*<0.001; inter-cluster: −58.95, *p*=0.001; CALM: intra-cluster=521.17, *p*<0.001; inter-cluster=21.10, *p*<0.001).

## Brain-defined subgroups differ in cognitive performance

Next, differences in cognitive performance between the clustering-defined groups were investigated. For the CALM sample, children who fell within the C1 range showed significantly higher performance across all cognitive assessments (all *p*_corrected_<0.05, see Figure 4A). For the ACE sample, C1 showed significantly lower performance across all cognitive measures, apart from visuospatial working memory (*p*=0.207, *p*_corrected_>0.999, see Figure 4B).

**Figure 4.**
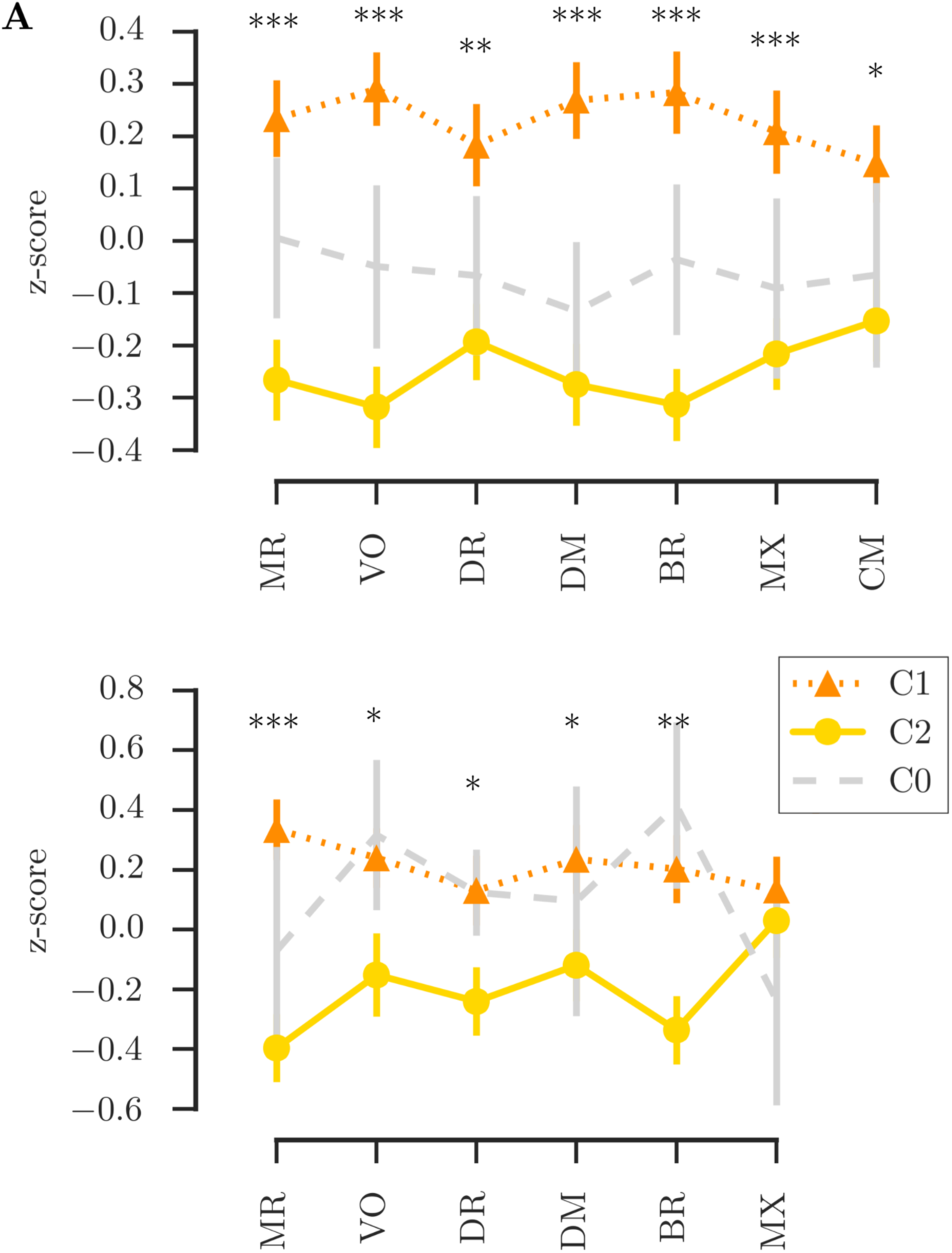
Comparison of cognitive scores between the groups defined based on cingulum FA in the CALM sample (**A**) and the ACE sample (**B**). Legend: Comparison between C1 and C2; C0: not assigned to C1 or C2, *** pcorrected<0.001, ** p_corrected_<0.01, * p_corrected_<0.05

We further investigated the reliability of clustering-defined grouping for predicting behavioural performance. For the CALM sample, predictive performance in unseen data was above chance for scores on Matrix Reasoning, Vocabulary, Mr. X, and Delayed Story Recall with prediction accuracy between 60 and 75% (see Table 2). For the ACE sample, there was above chance prediction for unseen data for Matrix Reasoning, Vocabulary, Backward Digit Recall, and Mr. X with prediction accuracies around 70% (see Table 2).

**Table 2.**
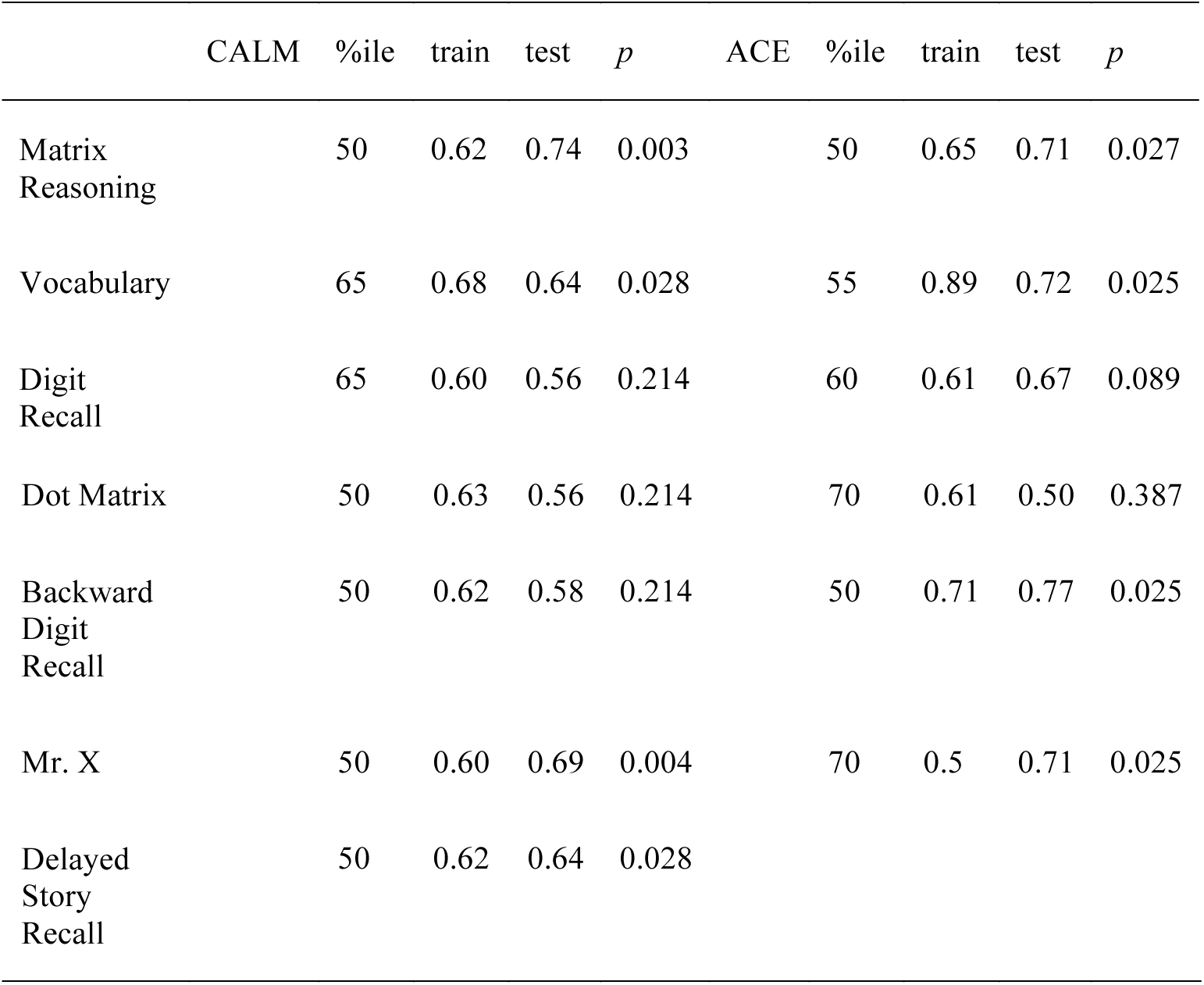
Generalizability of behavioural predictions based on clustering-defined groups. The optimal percentile cut-off identified in the training data set and the training performance are listed in the columns %ile and train. The test performance and statistical comparison to random assignment in the test set are listed in the columns test and *p*.

## Subgroups show differences in fronto-parietal and fronto-temporal connections of the cingulum

The groups were compared on the number of streamlines within the cingulum that connect cortical and subcortical regions. Streamlines associated with the cingulum were densest around in the core area of the cingulum, but also extended into the frontal, parietal, and temporal cortex in both hemispheres (see Figure 5A). Areas that were consistently connected through streamlines of the cingulum included the orbitofrontal and superior frontal cortex, insula, inferior and superior parietal cortex, precuneus, and parahippocampal cortex (see Figure 5B). The summed streamline density was higher in the left cingulum compared to the right cingulum (left: mean=53.34, SE=0.449; right: mean=45.80, SE=0.435, U=7959, *p*<0.001). The total number of connections was also higher in the left compared to the right cingulum (left: mean=44.10, SE=0.355; right: 35.31, SE=0.302, U=3282, *p*<0.001).Differences between groups in individual cingulum-mediated connections were selected using a cross-validation procedure, which indicated significant differences between the brain types for some cingulum connections that were also confirmed in a hold-out sample. The groups showed consistent differences in the connections between the precuneus and superiorfrontal cortex, inferior temporal cortex and lateral orbitofrontal cortex, parahippocampal cortex and post-central cortex in the left hemisphere, and the anterior cingulate cortex and posterior cingulate cortex in the right hemisphere (see Figure 5B). C2 had had lower streamline density in all of these connections compared to C1.

**Figure 5.**
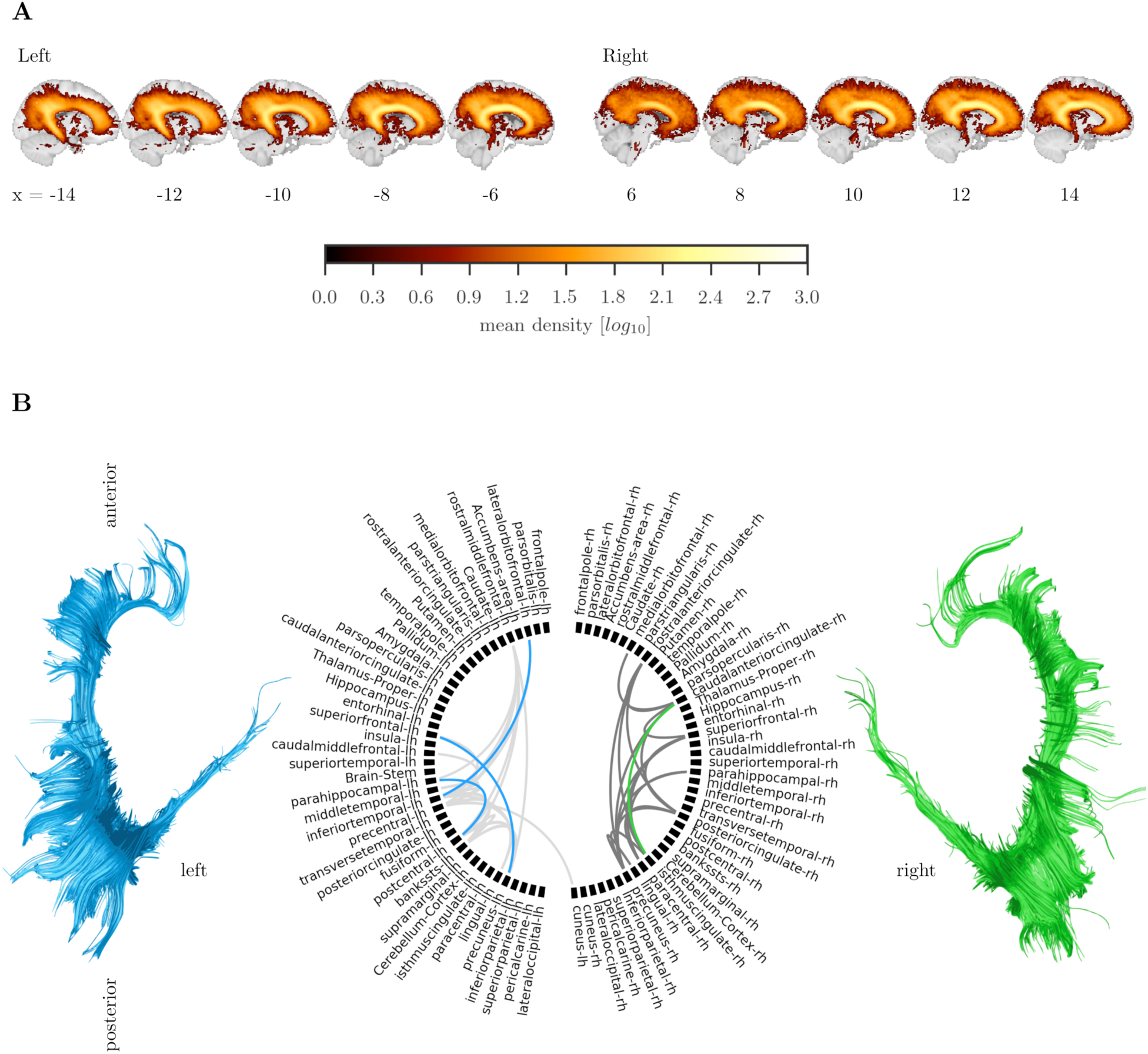
Structural connectivity of the cingulum. A: Average streamline density of the cingulum reconstruction. B: Connections of the cingulum. Grey lines indicate connections that were present in at least 60% of the sample. Blues lines indicate connections of the left cingulum that had fewer streamlines in group 2 compared to group 1 and green lines indicate fewer connections in group2 compared to group 1 for the right cingulum.

## Relationship between structural connectivity and default mode network activation

The default mode network (DMN) could be identified in group-level ICA (see Figure 6A). The groups did not differ in the number of components identified in individual independent component decomposition (C1: mean=100.70, SE=1.821; C2: mean=102.19, SE=1.862; U=1549, *p*=0.254). Comparison of ICA-derived maps between C1 and C2 indicated that the spatial extent of the DMN was significantly reduced in the posterior cingulate cortex for C1 (MNI peak coordinates: 0, −70, 35; TCFE-corrected *p*<0.05; see Figure 6B). Comparison of the DMN activation indicated reduced activation in C1 compared to C2 (C1: mean=-0.16, SE=0.137; C2: mean=0.13, SE=0.126, U=1372, *p*=0.049, see Figure 6D). Next, the relationship between variation in the microstructure of the cingulum and differences in functional connectivity of the medial prefrontal cortex (mPFC) and posterior cingulate cortex (PCC) was evaluated. While the clustering-defined groups did not differ in absolute mPFC-PCC partial correlation (C1: mean=0.31, SE=0.020; C2: mean=0.32, SE=0.020, U=1511, *p*=0.239), analysis of the relationship between cingulum FA and mPFC-PCC correlation indicated a significant group interaction (group x cingulum FA: *t*(118)=2.01, *p*=0.047). Follow-up analyses showed that cingulum FA was significantly associated with the mPFC-PCC correlation in C1 (beta=0.38, *p=*0.005), but not in C2 (beta=0.03, *p*=0.811, see Figure 6E).

**Figure 6A.**
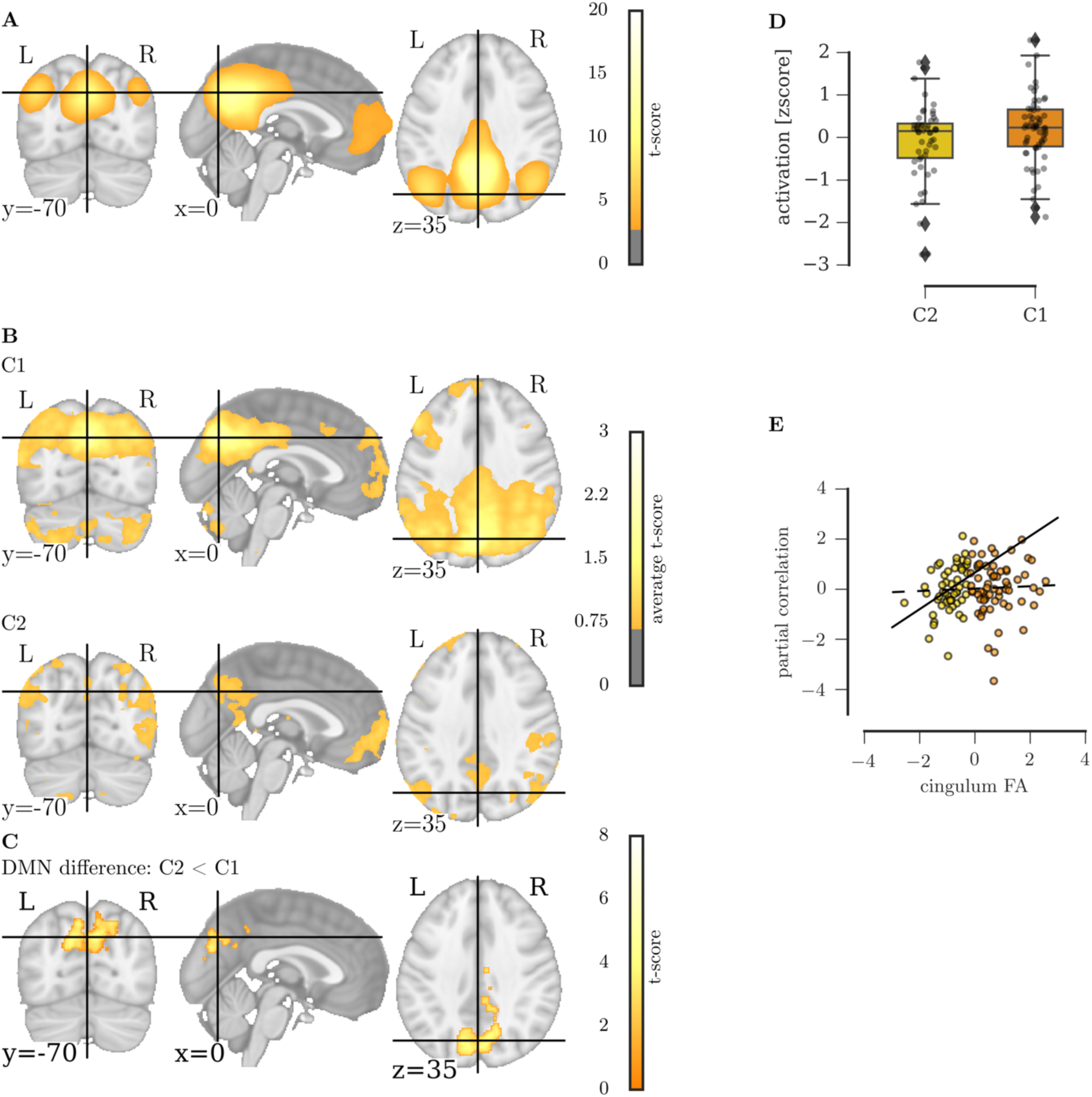
Default mode network (DMN) at the group level, **B** individual DMN average in C1 and C2, **C** Results of a two-sample t-test contrasting the DMN in C1 and C2 showing a reduced spatial extent of C1 compared to C2 in the posterior cingulate cortex. **D** DMN Activation in C1 and C2 **E** Relationship between the partial correlation of the medial prefrontal cortex and posterior cingulate cortex and FA of the cingulum in C1 and C2.

## Discussion

The first aim of the study was to identify groups that show individual differences in white matter organization. We employed data-driven community clustering to create subgroups of participants that were maximally similar in white matter organisation across 20 major white matter tracts. In theory, this could have resulted in multiple groups with a highly complex differentiation of brain organization. However, the subgrouping is dominated by the integrity of a pair of large tracts within the brain: the results indicated the presence of two groups primarily distinguished by FA of the left and right cingulum. Whilst this finding is somewhat unexpected, the cingulum has been implicated in a broad range of psychiatric and developmental disorders, including autism spectrum disorder (Ikuta et al. 2014), attention deficit hyperactivity disorder (Cooper, Thapar, and Jones 2015), schizophrenia (Abdul-Rahman, Qiu, and Sim 2011), major depression (Schermuly et al. 2010), mild cognitive impairment (Metzler-Baddeley et al. 2012), and dementia (Kantarci et al. 2011). One reason why such substantial and robust individual differences in the cingulum exist may be its prolonged development. The cingulum is one of the only tracts that show extended development with changes throughout childhood and adolescence, only reaching a stable level in the third decade of life (Lebel, Treit, and Beaulieu 2017; Tamnes et al. 2009). Similar tracts, also involved in long-range connectivity, like the superior longitudinal fasciculus, inferior longitudinal fasciculus, and inferior fronto-occipital fasciculus already plateau by the end of the second decade of life. Consequently, individual differences may arise from differences in timing of cingulum maturation. Alternatively, the cingulum may be particularly sensitive to environmental influences given that diffusion properties of white matter can change in response to environmental demands, e.g. in the context of motor or cognitive training (Caeyenberghs et al. 2016; Scholz et al. 2009). Inter-individual differences in cingulum FA could also, therefore, be a marker of accumulated differences in experience between individuals.

The second aim of the study of the study was to investigate whether subgroups formed on the basis of similarity of white-matter organization differ in cognitive performance. Children with lower cingulum FA showed lower performance on assessments of general intelligence, vocabulary, and short-term, long-term, and working memory. This broad difference in cognitive performance clashes with previous findings that indicated a comparatively narrow association between cingulum FA and executive function, which included sustained and divided attention (Takahashi et al. 2010), working memory (Golestani et al. 2014), and planning (Kubicki et al. 2003). There are several possible reasons for this contrast: First, the cognitive assessments in the current study were more comprehensive than in many previous studies. Second, the current study had much higher statistical power, while previous studies were likely to be underpowered with sample sizes ranging from 16 (Kubicki et al. 2003) to 38 participants (Takahashi et al. 2010).

Finally and most importantly, the effect observed here is likely larger and more widespread because individuals are *grouped on the basis of brain organization*. Grouping according to phenotype likely incorporates a high degree of heterogeneity in underlying neurobiology (Fair et al. 2012). The novel data-driven approach taken here created groups that maximise within-group homogeneity in neurobiology.

The prominent role of the cingulum in the brain-based grouping and the strong cognitive differences between the groups implies that this tract plays a critical role in the coordination of different brain regions. Indeed, the role of white matter structures is often seen as an extension of the brain regions that it connects (Catani et al. 2013). The cingulum contains strong connections between the medial prefrontal cortex (mPFC) and posterior cingulate cortex (PCC) and is related to their communication (Heuvel et al. 2008). We directly tested whether the functional connection between these regions is constrained by the cingulum. While a strong association between cingulum FA and mPFC-PCC was observed in a group of children with low cingulum FA, there was no association in children with high cingulum FA. A possible interpretation is that low cingulum FA limits the communication between these regions, but only up to a critical value. Once this critical value is reached – as in the case of the high FA group - further increases no longer influence the strength of the functional connection.

Integration between parietal and prefrontal areas is central to some theories of general intelligence (Deary, Penke, and Johnson 2010; Barbey 2017). According to the parietal-prefrontal integration theory (P-FIT), general cognitive ability arises from the interplay between parietal areas involved in the integration and abstraction of sensory information, prefrontal areas involved in reasoning and problem-solving, and the anterior cingulate cortex involved in response selection (Jung and Haier 2007; Barbey et al. 2012; Colom et al. 2009). The role of the cingulum may be to enable the efficient communication between these regions.

The connectivity mapping in the current study indicates that the cingulum plays a broader role in structural connectivity beyond connections along the prefrontal-parietal axis. The dense short-range and longer-range connections of the cingulum may enable communication within multiple large-scale structural brain networks. Indeed, the connections implicated in the group difference involved the prefrontal and posterior parietal areas that are part of a small number of so-called hub regions that are densely connected in functional and structural brain networks (Heuvel and Sporns 2011; Power et al. 2013). The connectivity of these hub regions is thought to form the backbone of a network architecture that enables the efficient transfer of information and the rapid transition between different network states, both of which may be necessary for cognitive performance in complex tasks (Barbey 2017).

There are some limitations to the current study. First, this study explored variation in white matter microstructure in pre-collected samples and was limited to the available data. Therefore, it was not possible to investigate if genetic or environmental factors contribute differentially to white matter microstructure in the brain types identified in the analysis. Further, while the brain types did not differ in age, there may be different underlying trajectories of brain development that result in differences in cingulum white matter microstructure at the time the participants were scanned. The influence of developmental trajectories can only be adequately resolved in longitudinal studies.

In conclusion, this is the first study to investigate the relationship between individual differences in white matter connections and cognitive performance with a brain-first approach. The results indicate that consistent individual differences exist in the microstructural organization of the left and right cingulum. Lower microstructural organisation of the cingulum was related to lower cognitive performance across a range of cognitive domains and was also linked to reduced BOLD activation within the default mode network. These findings suggest that cingulum connections play an important role in brain organisation and cognitive performance in childhood and adolescence. The current study is an initial step towards a bigger goal of understanding individual differences in neuroanatomy; it illustrates the benefits of bringing together large samples, multi-modal characterisation, and machine learning approaches like community clustering to identify biologically-defined groups that are difficult to distinguish at a behavioural level. In future, this new approach may enable our field to isolate key biological mechanism that drive differences in brain organisation, allowing for more objective identification of individual needs, and open new avenues for effective interventions.

## Acknowledgements

We want to thank Dr Edwin Dammaijer and Dr Rogier Kievit for helpful comments on early drafts of the paper. The Centre for Attention Learning and Memory (CALM) research clinic is based at and supported by funding from the MRC Cognition and Brain Sciences Unit, University of Cambridge. The Principal Investigators are Joni Holmes (Head of CALM), Susan Gathercole (Chair of CALM Management Committee), Duncan Astle, Tom Manly and Rogier Kievit. Data collection is assisted by a team of researchers and PhD students at the CBSU that includes Sarah Bishop, Annie Bryant, Sally Butterfield, Fanchea Daily, Laura Forde, Erin Hawkins, Sinead O’Brien, Cliodhna O’Leary, Joseph Rennie, and Mengya Zhang. The authors wish to thank the many professionals working in children’s services in the South-East and East of England for their support, and to the children and their families for giving up their time to visit the clinic.

## Conflicts of interest

The authors declare no conflict of interest

